# Structural properties and peptide ligand binding of the capsid homology domains of human Arc

**DOI:** 10.1101/2020.12.17.423319

**Authors:** Erik I. Hallin, Clive R. Bramham, Petri Kursula

## Abstract

The activity-regulated cytoskeleton-associated protein (Arc) is important for synaptic scaling and the normal function of the brain. Arc interacts with many neuronal postsynaptic proteins, but the mechanistic details of its function have not been fully established. The C-terminal domain of Arc consists of tandem domains, termed the N- and C-lobe. The N-lobe harbours a peptide binding site, able to bind to multiple targets. By measuring the affinity of various peptides towards human Arc, we have refined the specificity determinants of this site. We found two sites in the GKAP repeat region that may bind to Arc and confirmed these interactions by X-ray crystallography. Comparison of the crystal structures of three human Arc-peptide complexes identifies 3 conserved C-H...*π* interactions at the binding cavity, which explain the sequence specificity of short linear motif binding by Arc. By analysing the structures, we further characterise central residues of the Arc lobe fold, show the effects of peptide binding on protein dynamics, and identify acyl carrier proteins as structures similar to the Arc lobes. We hypothesise that Arc may affect protein-protein interactions and phase separation at the postsynaptic density, affecting protein turnover and re-modelling of the synapse.

## Introduction

Normal brain function depends on signal transduction across synapses, and the ability to modify the strength of synaptic transmission is key to learning and memory. Synaptic transmission and plasticity involve many proteins located on both the pre- and postsynaptic side of the synapse. One of the central proteins in long-term synaptic plasticity is the activity-regulated cytoskeleton-regulated protein (Arc) (1, 2). Arc is an immediate early gene expressed in excitatory, glutamatergic neurons following synaptic activation. Arc mRNA transports into dendrites, where it undergoes local translation near synapses. In neuronal activity-dependent synaptic plasticity, such as long-term potentiation (LIP) and long-term depression (LTD), Arc protein is rapidly expressed and degraded, indicating a transient and dynamic mode of action (1, 3–6). Transient expression of Arc is also required for memory consolidation and postnatal developmental plasticity of the visual cortex (7–10).

Arc is a flexible hub, interacting with many different proteins located in the postsynaptic density and the nucleus (11). The abundance of AMPA-type glutamate receptors (AMPARs) at the synapse is a major determinant of synaptic transmission efficiency. Expression of Arc promotes endocytosis of AMPARs, by recruitment of the machinery for clathrin-mediated endocytosis, resulting in synaptic depression (12–14). Recently, Arc was found to bind stargazin (Stg), an auxiliary subunit of AMPAR important for receptor trafficking (15–17). Arc is also able to form capsid-like structures and transfer nucleic acids between neurons (18–21). The C-terminal domain of Arc (Arc-CT) consists of two structural repeats called the N-lobe (Arc-NL) and the C-lobe (Arc-CL). These domains are structural homologs to retroviral capsid proteins, but the N-lobe domain differs by having a peptide binding site, binding to stargazin and other peptides (16, 22).

The peptide binding site of the Arc-NL allows an extended peptide to bind across a hydrophobic pocket. Together with the N-terminal tail of Arc-NL, which is located on top of the peptide in a parallel manner with the peptide, both parts are forming a short beta-sheet. The protein-peptide interaction does not seem to be highly specific since the protein barely interacts with the peptide sidechains and mostly interacts with the peptide backbone (23). Based on alignments of Arc ligand peptides, the PxF/Y/H binding motif was proposed (16). The histidine of the motif became questionable when we in an earlier study showed that the GluN2A peptide, having PxH and suggested to bind Arc in the peptide binding pocket (16), did not bind to hArc-NL, using the more robust method ITC (23). Further characterisation of the specificity determinants of the Arc-NL peptide binding site would be of high value for finding novel Arc binding partners.

The postsynaptic density (PSD) is a protein-rich assembly below the postsynaptic membrane, formed of large scaffolding proteins. These proteins carry a combination of protein interaction domains, which may interact with several alternative partners; the structure of the protein assembly can be regulated in an activity-dependent manner. The main PSD scaffolds include membrane-associated guanylate kinases (such as PSD-95), Shanks, calmodulin-dependent protein kinase II, Homer, synaptic Ras GTPase-activating protein 1, and guanylate kinase-associated protein (GKAP). Scaffolding proteins of the PSD are targets for mutations in neurological disorders (24). GKAP is a structural protein of the PSD, linking PSD95 to Shank, which accumulates during synaptic inactivity (25). The binding site of GKAP for Shank is located at the C terminus of GKAP, interacting with the Shank PDZ domain (26–28). GKAP has 5 sequence repeats located in its middle region, interacting with PSD95 (29, 30), and one repeat overlaps with the previously suggested Arc binding site (16, 23). PSD95 is required for targeting of Arc to the PSD, where it is found in multiple PSD95 interaction complexes (31). It is therefore possible that Arc expression contributes to activity-dependent remodelling of the PSD.

To gain insight to human Arc signalling, we sought to characterise the structural determinants of ligand binding to the hArc-NL. We analysed the specificity of peptide binding by human Arc to find out the required residues for binding. By measuring the affinity between hArc-NL and various peptides, we show that the peptide binding site is sequence-specific. Mutation of the proline or the tyrosine of the PxF/Y motif resulted in loss of binding. We found two Arc interaction sites on GKAP, suggesting that Arc may bind to the repeat region of GKAP. By X-ray crystallography, we confirm the peptide binding on hArc-NL and provide the structure of hArc-CL, which does not have the peptide binding ability found in hArc-NL.

## Material and methods

### Recombinant protein production

The expression and purification of hArc NL and CL were performed as described (23). In short, hArc-NL (residues 207-277), hArc-CL (residues 277-370), and hArc-CT (residues 206-396, containing both lobes) were expressed with a cleavable His-MBP tag in *E. coli* BL21, with isopropyl [β3-D-1-thiogalactopyranoside induction at +30 °C. The cells were lysed by sonication and the proteins purified by Ni-NTA affinity chromatography and size-exclusion chromatography. The affinity tag was removed with recombinant TEV protease during purification.

### Peptide synthesis

Peptides were synthesised by GenScript (Piscataway NJ, USA), carrying N-terminal acetylation and C-terminal amidation. The sequences and nomenclature of the synthetic peptides are given in Table 1.

**Table 1.**
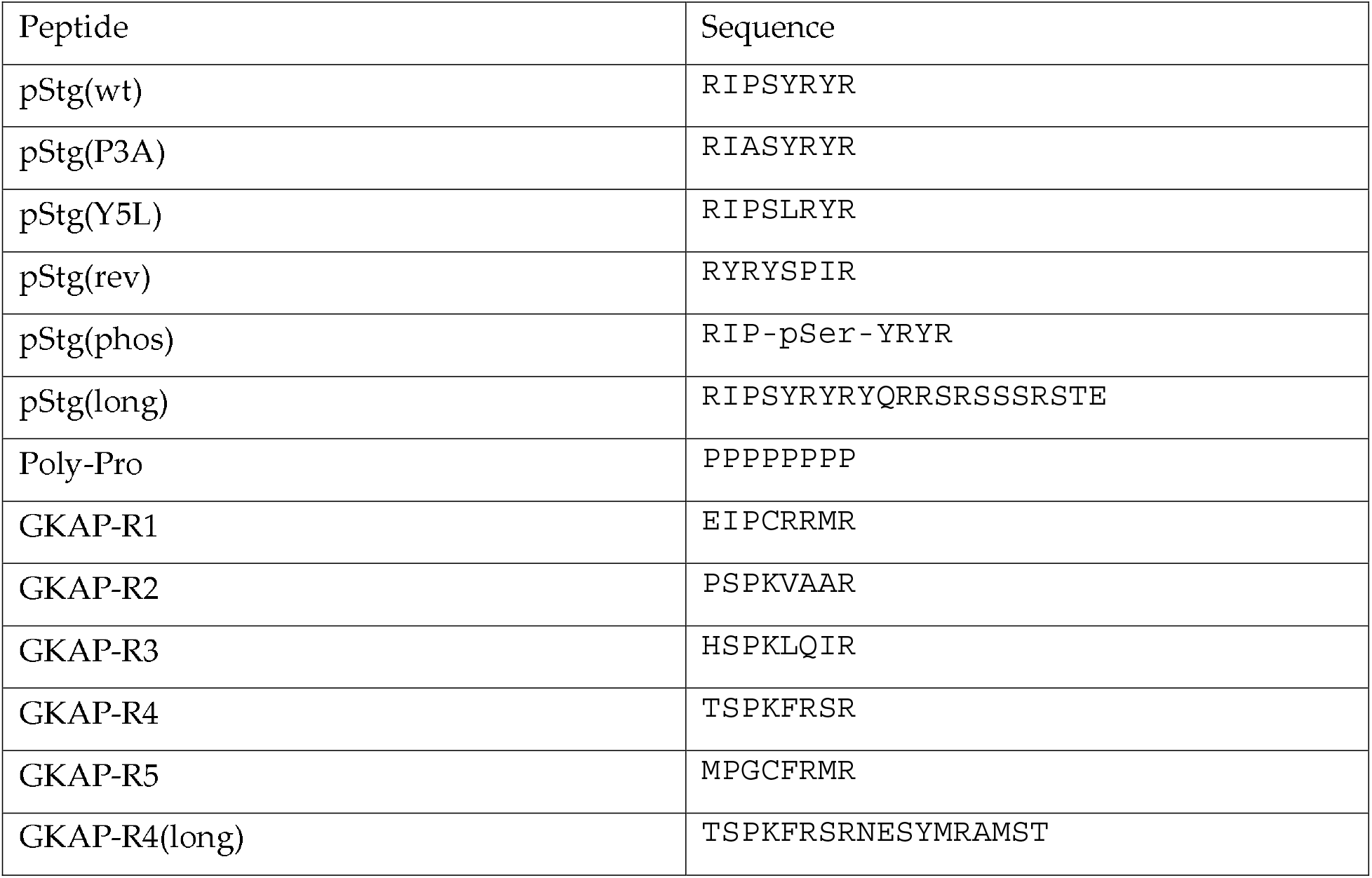
Synthetic peptides used in the experiments.

### Isothermal titration calorimetry

The affinity of peptides to hArc lobe domains was measured using a MicroCal iTC200 (Malvern, UK) instrument. The peptide was injected into the cell containing the purified protein; the peptides were 2.5-5.0 mM and the protein 0.25-0.5 mM. The buffer used was 20 mM Tris-HCl, pH 7.4, with 150 mM NaCl, and the measurements were performed at +25 °C. The data were analysed using MicroCal Origin 7, using a one-site binding model.

### Protein crystallisation

Crystals of hArc lobes were obtained by vapour diffusion. The proteins and peptides were in a buffer consisting of 20 mM of Tris-HCl and 150 mM of NaCl. The crystal of hArc-NL with pStg(wt) was made by mixing hArc-NL (20 mg/ml) with a 3-fold molar excess of pStg(wt). Hanging drops were set up by mixing 1 μl of the protein/peptide solution with 1 μl of the reservoir, consisting of 1.4 M trisodium citrate, and incubated at +4 °C. The crystals were soaked in PEG400 as cryo-protectant for a few seconds before flash-cooling in liquid nitrogen. The crystal of hArc-NL together with GKAP-R4 was made by mixing hArc-NL (20 mg/ml) with a 3-fold molar excess of the peptide. Hanging drops were made by mixing 2 μl of the protein/peptide mixture with 2 μl of the reservoir, consisting of 1.2 M of trisodium citrate, and incubated at +20 °C. A solution of 1.2 M trisodium citrate with 20% glycerol was used as cryoprotectant for these crystals. The crystal of hArc-NL with GKAP-R5 was made by mixing hArc-NL (20 mg/ml) with a 2-fold molar excess of peptide. Hanging drops were made by mixing 2 μl of the protein/peptide mixture with the reservoir, consisting of 1.2 M of trisodium citrate, and incubated at +4 °C. Crystals were picked and flash-frozen in liquid nitrogen. Crystals of hArc-CL were obtained by sitting-drop vapour diffusion. The drops were made by mixing 200 nl of the protein (15 mg/ml) with 100 nl of a reservoir, consisting of 16% PEG 8000, 40 mM potassium-phosphate (monobasic), and 20% glycerol. The drops were incubated at +20 °C, and crystals were picked and flash-frozen in liquid nitrogen.

### X-ray diffraction data collection and structure refinement

Diffraction data for hArc-NL with pStg(wt) were collected on the 103 beamline at Diamond Light Source (Oxfordshire, UK). Data for hArc-NL with GKAP-R4 and GKAP-R5 were collected on the P13 beamline (32) at EMBL/DESY (Hamburg, Germany). Data for hArc-CL were collected on the P14 beamline at EMBL/DESY (Hamburg, Germany). All data were collected at 100 K and processed using XDS (33). Phasing was done by molecular replacement, using PDB entry 4×3H (16) for hArc-NL and PDB entry 4X3X (16) for hArc-CL as search models in Phaser (34). The structures were refined using phenix.refine (35) and model building was done in Coot (36). Structure validation was done using MolProbity (37).

### Structure analyses

Network analyses on the protein structures were carried out on the NAPS server (38, 39). Structure similarity searches were done using Salami (40). The pStg(phos) peptide was modelled into hArc-NL manually in Coot, based on the crystal structure, and energy minimisation was carried out in Yasara (41). Molecular dynamics (MD) simulations for hArc-NL without peptide, and with either pStg(wt) or pStg(phos) bound, were run with Yasara, essentially as described (42). The simulations were run for 200-250 ns at +25 °C, and trajectory analysis was carried out in Yasara.

## Results and discussion

We set out to study the structure of the two lobes of Arc-CT in the human protein, and to better understand the binding of short linear peptide motifs to hARc-NL. We further wanted to observe possible peptide interactions of hArc-CL and the effects of Stg phosphorylation on the interaction with hArc-NL. Using a combination of ITC and X-ray crystallography, we provide an improved understanding of hArc structure and refine the linear peptide motif that interacts directly with the binding site on hArc-NL.

### Peptide ligand binding by the human Arc lobe domains

To study the specificity of peptide binding by hArc-NL, we measured the affinity of a set of mutated versions of a known peptide ligand from Stg. The Stg peptide has shown the highest affinity to hArc-NL in our earlier studies (23) and has the suggested PxF/Y motif (16). For clarity, we number the binding motif such that the conserved aromatic residue is P(0). Our results show that by mutating the consensus sequence, affinity to hArc-NL is lost (Fig. 1A, Table 2). As controls, hArc-NL did not bind to a poly-proline peptide nor to the reverse sequence of Stg. The modifications of P3A and Y5L also resulted in loss of binding, showing that the binding site is specific and that the proline and tyrosine are important for binding to hArc-NL. The binding of pStg(wt) to hArc-NL was confirmed by X-ray crystallography of the complex (see below), and the binding mode is similar to the earlier structure of the same complex at lower resolution (16). Crystallisation screening of hArc-NL with the mutated peptides was unsuccessful, which is an additional indication that hArc-NL does not bind these peptides, since Arc-NL does not seem to crystallise without a bound peptide.

**Figure 1.**
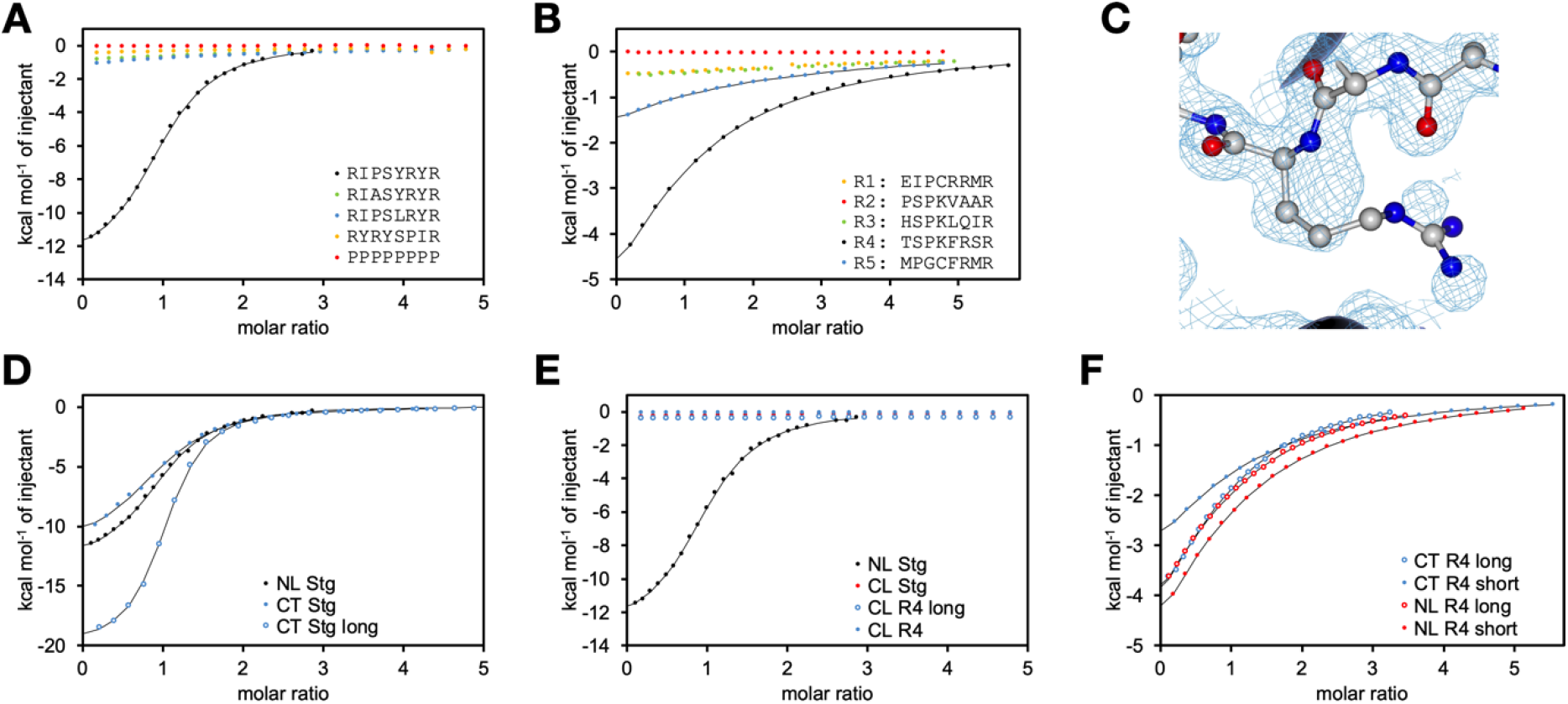
Calorimetric analysis of hArc ligand peptide binding. A. Titration of hArc-NL with pStg variants. B. Titration of hArc-NL with GKAP repeat peptides. C. Electron density for structure 4×3i (16), which clearly has electron density for Pro instead of Arg in the ligand peptide. D. Comparison of hArc-NL and hArc-CT binding to Stg. E. Comparison of hArc-NL and hArc-CL. F. Comparison of short and long GKAP-R4 peptide binding to hArc-NL and hArc-CT.

**Table 2.** ITC data for Arc-peptide interactions. Only combinations of Arc domain and peptide with detectable binding in ITC are shown (see Fig. 1 for all binding curves). The hArc-CL construct did not bind to any peptide, and repeats 1-3 of GKAP did not show detectable binding to any construct in the experiments.

| Protein | Peptide | K<sub>d</sub> (μM) | ΔH (kcal/mol) | -TΔS (kcal/mol/°) |
| --- | --- | --- | --- | --- |
| hArc-NL | pStg(wt) | 33.6 ± 1.1 | -13.1 ± 0.1 | +7.0 |
| hArc-NL | pStg(phos) | 23.8 ± 1.4 | -66.5 ± 0.6 | +60.2 |
| hArc-NL | GKAP-R4 | 714 ± 19 | -10.9 ± 0.4 | +6.6 |
| hArc-NL | GKAP-R5 | 2390 ± 314 | -8.2 ± 0.3 | +4.7 |
| hArc-NL | GKAP-R4(long) | 426 ± 12 | -7.2 ± 0.2 | +3.1 |
| hArc-CT | GKAP-R4 | 763 ± 10 | -6.8 ± 0.1 | +2.5 |
| hArc-CT | GKAP-R4(long) | 391 ± 18 | -6.8 ± 0.2 | +2.2 |
| hArc-CT | pStg | 80 ± 4.0 | -12.0 ± 0.2 | +6.4 |
| hArc-CT | pStg(long) | 26.4 ± 1.2 | -20.2 ± 0.2 | +14.0 |

hArc-NL binding to GKAP repeat peptides was tested next, as one of these repeats was shown to be a ligand previously (16, 23). The affinity of hArc-NL towards the five repeat sequences of GKAP showed binding in two cases (Fig. 1B, Table 2). GKAP-R4, which was known to bind (16, 23), showed higher affinity. GKAP-R5 also bound to hArc-NL, while the other repeats did not show measurable affinity using ITC (Fig. 1B). The binding of GKAP-R4 and GKAP-R5 was confirmed with X-ray crystallography, and the peptides were bound in the same peptide binding site of hArc-NL (see below). The selective binding of these two repeats reveal new information on hArc binding specificity. Both GKAP-R4 and GKAP-R5 have the phenylalanine of the PxF/Y motif, while the other sequences have Arg, Val, or Leu at this position. All repeats except for GKAP-R5 have the proline of the PxF/Y motif. Still, GKAP-R5 binds to hArc-NL. It has a neighbouring Pro residue one position before the canonical Pro, which is replaced by a Gly. It is likely that the conserved Pro residue is important for both keeping an extended conformation for the binding peptide and in making specific interactions, such as C-H...π bonds towards Arc-NL. The extra Gly residue in GKAP-R5 provides needed flexibility to accommodate this altered motif into the binding site. Overall, the results show that both the Pro and Phe/Tyr residue of the motif are required for short peptide ligand binding to hArc-NL, but the position is not absolutely fixed.

Using fluorescence polarisation spectroscopy, two other peptides were reported to bind to Arc-NL, which both have His instead of the Phe/Tyr residue of the PxF/Y motif (16). Using ITC in our earlier work (23), we could not see any indication of Arc-NL binding to the GluN2A peptide. Indeed, Arc binds to another segment of GluN2A having a PxF/Y motif (22). These observations indicate that the Phe/Tyr residue is important for binding.

An unconventional CaMKII peptide was reported to bind to Arc-NL, in which the canonical Pro is replaced by Arg (16). However, observation of the electron density of the corresponding crystal structure reveals that the peptide sequence in the structure is erroneous, as the density indeed shows a peptide with a Pro at the canonical position instead of Arg (Fig. 1C). This is a further indication of the sequence specificity of the binding pocket in Arc-NL.

Arc-CL is a structural homologue of Arc-NL, and it could in theory have a similar ligandbinding site. It might be possible that a long peptide that binds to Arc-NL would continue into the corresponding region of Arc-CL, forming an extended peptide binding surface. However, we could not detect binding of hArc-CL to any of the tested peptides (Fig. 1D). Therefore, we tested longer versions of Stg and GKAP peptides that bind hArc-NL to see if Arc-CT, with both lobes present, has altered affinity towards them. However, the affinity towards the short and long peptides was comparable (Fig. 1E-F, Table 2), suggesting that hArc-CL did not interact with the C-terminal region of the longer peptide. A similar small increase in affinity could be seen towards the longer peptides for both hArc-NL alone and hArc-CT (Table 2), showing that hArc-CL is not involved. For the longer Stg peptide, hArc-CT bound with a higher favourable enthalpy than hArc-NL; this could be an indication of additional interactions between the peptide and hArc-CL. However, the change in affinity was small, and the binding entropy was more unfavourable, compensating for the enthalpy contribution.

The NMR structure of the rat Arc-CT (22), consisting of both lobes, showed that the relative orientation of the two lobes could allow a peptide binding to the Arc-NL to continue into the Arc-CL if extended. However, it appears that the peptide binding feature is unique to the mammalian Arc-NL. The structure of *Drosophila* Arc1 without bound peptides showed that the sequence upstream of the NL could fold into the peptide binding pocket (43). This folding back would block peptide binding and dArc may be unable to bind to peptide ligands like mammalian Arc-NL. The structures of dArc capsids (44) suggest the same. The Stg peptide that binds to human and rat Arc-NL does not bind to the two isoforms of drosophila Arc (45).

All peptides showed similar enthalpy-entropy compensation, when binding thermodynamics were assessed using ITC (Table 2). The observed thermodynamics are typical for a proteinligand interaction in the micromolar affinity range (46, 47). Favourable enthalpy is brought about by hydrogen bonds as well as weaker interactions including C-H...π bonds and van der Waals interactions, while unfavourable entropy can be explained by overall loss of flexibility in the complex compared to molecules free in solution.

### Structures of hArc-NL bound to ligand peptides

We aimed to obtain more detailed information on Arc-peptide interactions using X-ray crystallography. A comparison of the crystal structures of the different Arc-peptide complexes (Table 3) further highlights important details of peptide binding (Fig. 2). The structures of rat Arc-NL and Arc-CL were previously reported (16). We report the hArc-NL structure bound to the pStg(wt) peptide at higher resolution (1.90 Å) (Fig. 2A). The structure of hArc-NL bound to GKAP-R4 shows that the peptide backbone and the PxF motif align well with pStg(wt) (Fig. 2B). hArc-NL complexed with GKAP-R5 is different, showing that the peptide backbone is shifted at the position of the canonical Pro at P(−2), so that the neighbouring Pro at P(−3) present in GKAP-R5 can get closer to the binding pocket (Fig. 2C). Thus, the linear motif for Arc-NL binding can be expanded from from PxF/Y to P(G)xF/Y.

**Figure 2.**
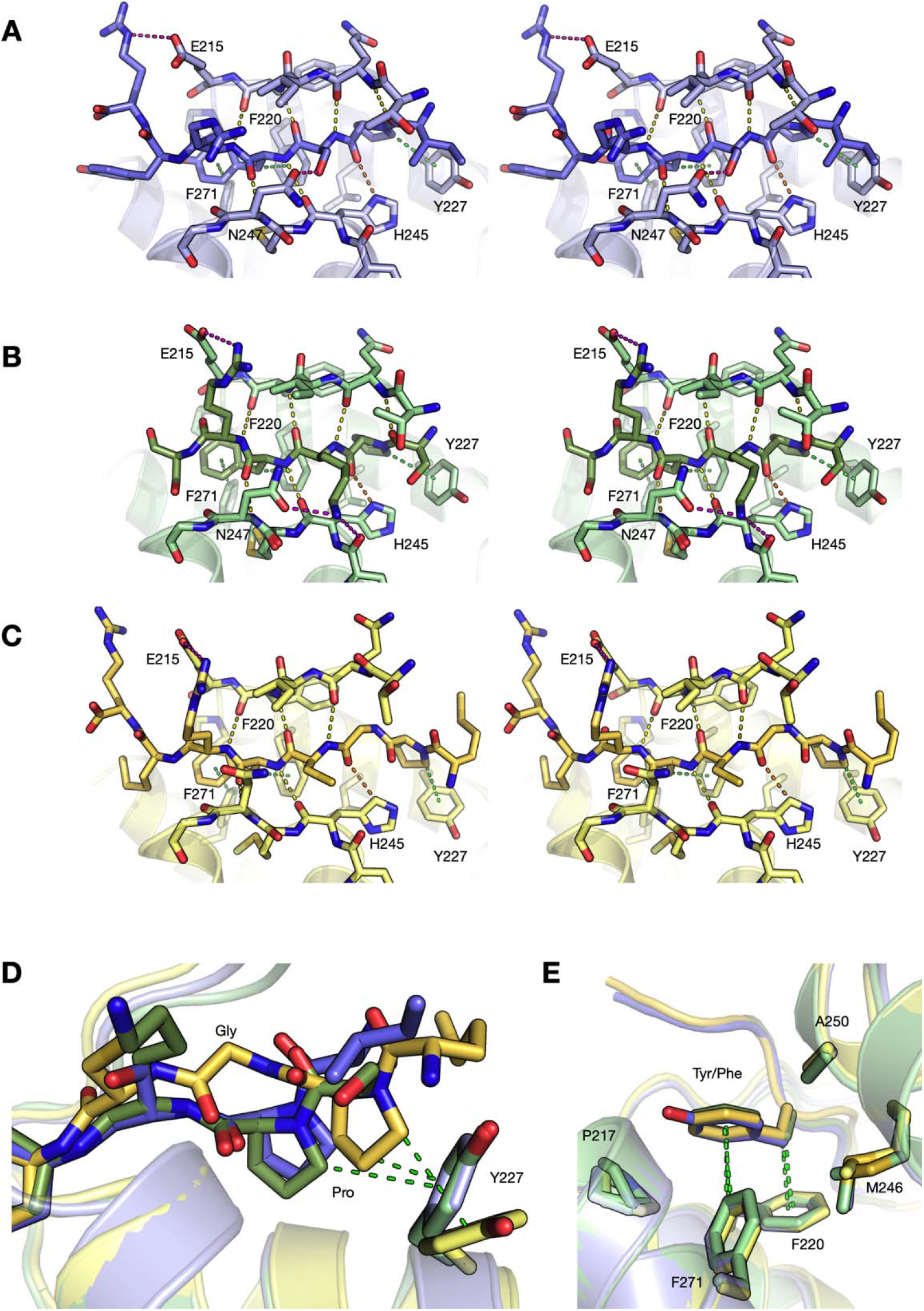
Structures of hArc-NL complexes with peptide ligands. A. Stereo view of the hArc-NL complexed with pStg(wt). B. Stereo view of the hArc-NL complexed with GKAP-R4. C. Stereo view of the hArc-NL complexed with GKAP-R5. D. Conserved C-H...π interactions of the Pro residue in the ligand peptie with Tyr227. E. Two conserved C-H...π interactions of the peptide aromatic residue.

**Table 3.** Crystallographic data processing and refinement.

| Structure | hArc-NL +<br>pStg(wt) | hArc-NL +<br>GKAP-R4 | hArc-NL +<br>GKAP-R5 | hArc-CL |
| --- | --- | --- | --- | --- |
| PDB entry | 6TNO | 6TNQ | 6TQ0 | 6TN7 |
| Beamline | P13 | P13 | P13 | P14 |
| Wavelength | 0.976 | 0.976 | 0.976 | 0.976 |
| Space group | P2<sub>1</sub> | P2<sub>1</sub> | P2<sub>1</sub> | P22<sub>1</sub>2<sub>1</sub> |
| Unit cell<br>parameters | 51.39 29.77 71.79<br>90 92.5 90 | 50.80 29.37 71.36<br>90 93.4 90 | 39.31 192.30<br>39.56 90 94.8 90 | 37.21 38.29 61.01<br>90 90 90 |
| Resolution range<br>(Å) | 50-1.90 (1.95-1.90) | 50-1.30 (1.34-1.30) | 50-1.95 (2.00-1.95) | 50-1.67 (1.71-1.67) |
| Completeness<br>(%) | 97.8 (97.9) | 89.5 (40.4) | 98.2 (97.3) | 92.2 (32.5) |
| R<sub>sym</sub> (%) | 4.1 (38.6) | 2.7 (13.8) | 4.5 (178.6) | 5.1 (62.6) |
| R<sub>meas</sub> (%) | 4.8 (45.1) | 3.2 (17.9) | 5.3 (207.3) | 5.5 (69.9) |
| CC<sub>1/2</sub> (%) | 99.9 (92.1) | 99.9 (96.3) | 99.9 (40.0) | 99.9 (85.7) |
| <I/σ(I)> | 18.6 (2.9) | 24.2 (4.6) | 13.8 (0.9) | 18.3 (2.6) |
| Redundancy | 3.6 (3.7) | 3.4 (2.0) | 3.7 (3.9) | 7.1 (4.5) |
| Wilson B (Å<sup>2</sup>) | 36.3 | 22.9 | 53.8 | 35.2 |
| R<sub>work</sub>/R<sub>free</sub> (%) | 18.2/21.6 | 13.2/17.0 | 22.1/26.3 | 24.2/26.8 |
| Rmsd bond<br>length (Å) | 0.005 | 0.016 | 0.012 | 0.004 |
| Rmsd bond<br>angle (°) | 0.9 | 1.4 | 0.8 | 0.8 |
| Ramachandran<br>favoured/outliers<br>(%) | 100/0 | 99.5/0 | 99.5/0.2 | 98.8/0 |
| MolProbity score<br>(percentile) | 1.74 (86<sup>th</sup>) | 1.68 (61<sup>st</sup>) | 1.62 (92<sup>nd</sup>) | 1.72 (77<sup>th</sup>) |

The key interactions in all complexes include three C-H...π hydrogen bonds that directly involve the P(G)xF/Y motif. Pro at P(−2/-3) forms a C-H...π interaction with the aromatic ring of Tyr227 in Arc (Fig. 2D). In addition, Tyr/Phe(0) is involved in two C?.,.π interactions (Fig. 2E). The C(3 atom interacts with the side chain of Phe220, and the aromatic ring accepts a C-H...π bond from Phe271 of Arc. Other residues in the peptides also make conserved interactions with hArc-NL (Fig. 2). The visually strongest of these are main-chain [β3-sheet interactions of the bound peptide on both sides of the chain. Additionally, the conserved Arg at P(+1) and/or P(+3) form salt bridges to Glu215 of hArc-NL. Asn247 interacts with the side chain of the variable residue in the P(−1) position in all the three complexes, which may be one affinity determinant between different ligand motifs.

The crystal structure of hArc-NL was subjected to MD simulations, with and without the pStg(wt) ligand. This was done in order to follow the dynamics of the Arc-NL N-terminal region that in peptide complexes, folds as a β strand on top of the peptide. The results indicate flexibility of the N terminus of hArc-NL in the absence of a peptide ligand (Fig. 3A). Furthermore, the radius of gyration during simulation is smaller for the peptide complex than the free protein (Fig. 3B-C). Hence, conformational changes in Arc may be linked to peptide ligand binding, as the flexible linker lies N-terminal to hArc-NL.

**Figure 3.**
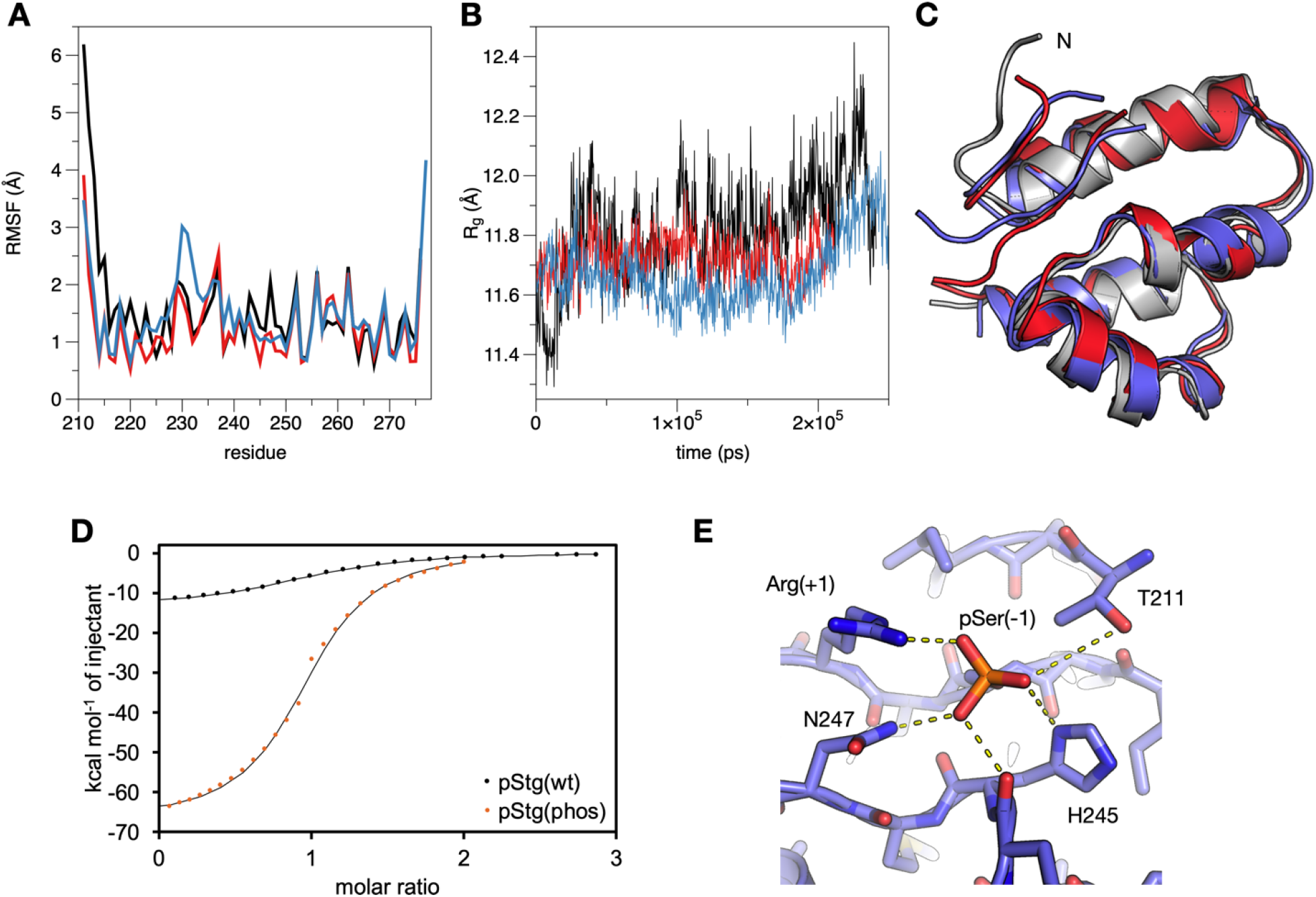
Simulations and Stg phosphorylation. A. Root mean-square fluctuations (RMSF) of hArc-NL residues without the peptide (black), with pStg(wt) (red), and with pStg(phos) (blue). Note the flexible N terminus in the absence of ligand peptides and the increased mobility of helix 2 in the presence of pStg(phos). B. Radius of gyration during the simulations; colouring as in (A). C. Structural snapshots from each simulation at a time point of 200 ns. D. ITC analysis of hArc-NL binding to pStg(phos). E. Polar interactions of phospho-Ser228 200 ns into the simulation.

### Phosphorylation of Stargazin

Arc interacts with Stg through the binding site on Arc-NL, and Stg phosphorylation state is regulated during synaptic plasticity (48). One phosphorylation site is located in the middle of the Stg peptide, at the P(−1) position of the binding motif PxF/Y. We tested if phosphorylation of the Stg peptide at Ser228 affected the binding affinity to hArc-NL. The binding entropy and enthalpy both changed, but binding affinity remained the same (Fig. 3D, Table 2), indicating that the Stg motif binds to hArc-NL regardless of the phosphorylation state at Ser228. Our result is in conflict with an earlier experiment, where a fluorescence polarisation competition binding assay showed that phosphorylation of a Stg peptide at Ser228 reduced the affinity to Arc-NL 20-fold (16). Like for the other studied peptides, a strong enthalpy-entropy compensation is present (Table 2), and the observed changes are in line with those expected, when comparing peptides with and without phosphorylation (49). The high favourable enthalpy is a sign of electrostatic interactions between the phosphate group and the protein, while the large unfavourable entropy component is likely related to the more flexible nature of the phosphopeptide when free in solution, as well as less flexibility of the Arc partner in the pStg(phos) complex.

The structure of hArc-NL bound to pStg(wt) shows that Ser228 points away from the hydrophobic pocket, and phosphorylation should not generate major clashes. A model of hArc-NL bound to pStg(phos) was made based on the crystal structure to get an insight into the possible additional interactions. The phospho-Ser228 side chain is surrounded by potentially interacting residues from all sides (Fig. 3E). The residues most likely making contact with the phosphate group include Asn247, His245, and Thr211. MD simulations further indicate that the Arg side chain two residues downstream on the peptide can make a salt bridge contact with the phospho-Ser228; this explains some of the favourable enthalpy and unfavourable entropy observed in ITC. At a more coarse level, it can be seen that the phosphopeptide binding is accompanied by an increased flexibility of the first half of helix ፧2 of hArc-NL (Fig. 3A).

### Structure of hArc-CL and folding properties of the lobes

We crystallised hArc-CL, which has the same overall fold as hArc-NL (Fig. 4A). As expected, hArc-CL is similar to rat Arc-CL, but gives a higher resolution and shows the structure of the missing loop after the first helix (Fig. 4B). The largest variation between the rat and human Arc-CL is the tilt of the first helix (here termed ፧0), which is a sign of flexibility of the structure, and could even reflect varying conformations of the Arc-CT in the context of full-length Arc. The sequences of rat and human Arc-CL only differ by 3 residues, and the structures are expected to be similar. One of these residue variations is Phe333 in rat Arc, which is Leu for hArc. The smaller residue in hArc allows the second helix to come closer to the protein core. However, whether this is of functional relevance, remains to be studied.

**Figure 4.**
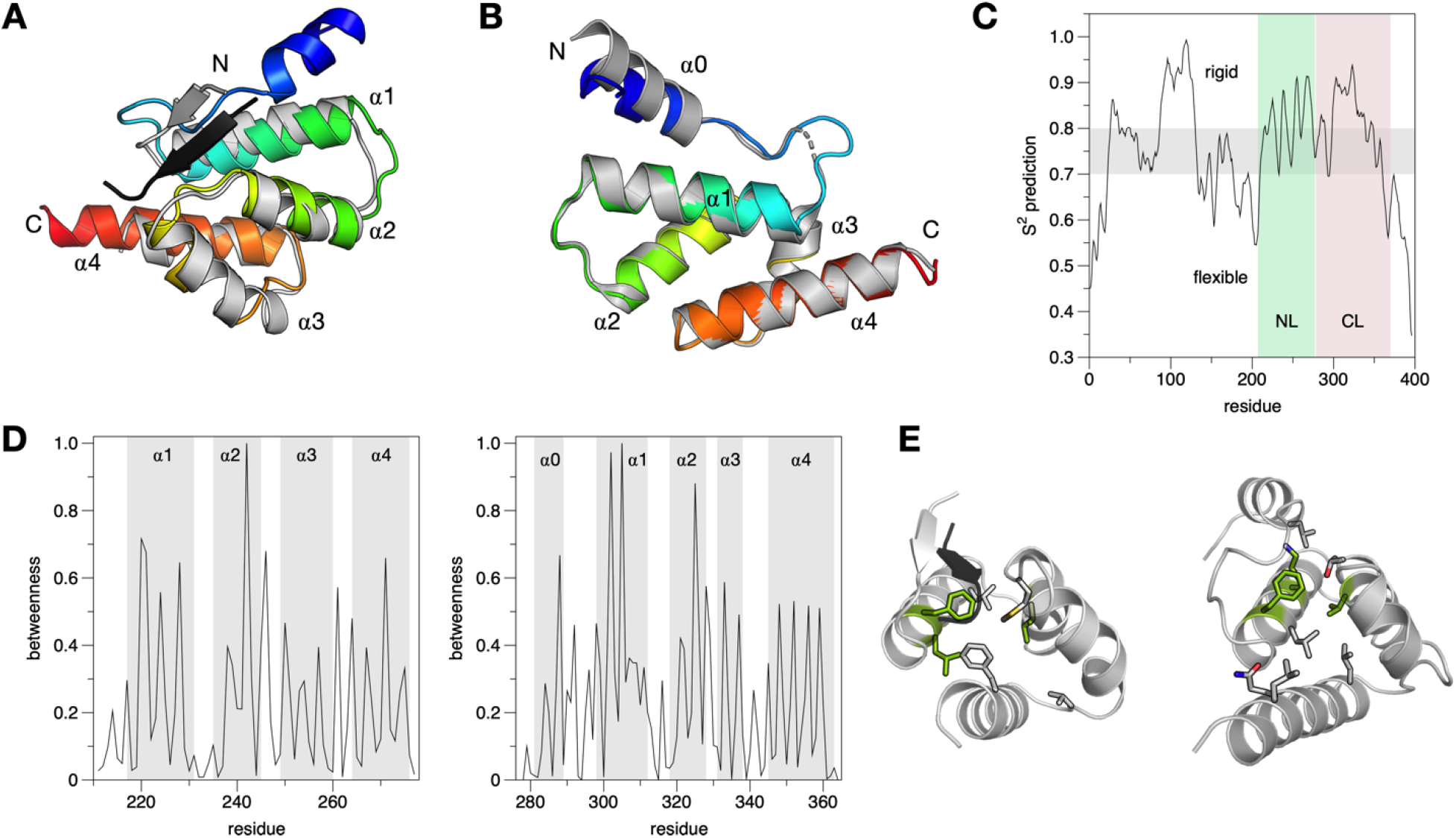
Structure of hArc-CL and structure analyses. A. Superposition of hArc-NL (grey) and hArc-CL (rainbow colours). B. Superposition of human (coloured) and rat (grey) Arc-CL. C. DynaMine analysis of human full-length Arc. D. Betweenness analysis of hArc-NL (left) and hArc-CL (right). E. 3D view of the most central residues, based on the betweenness scores. Sidechains of residues scoring >0.5 are shown, and the top 3 central residues in each domain are coloured green. Left: hArc-NL, right: hArc-CL.

A sequence-based prediction of chain dynamics was carried out for full-length Arc (Fig. 4C). The most rigid predicted region is the oligomerisation region (19) in the helical coil of Arc-NT. The Arc-CT helices are also predicted as rigid, while the termini and the central linker are flexible. All these aspects comply well with current experimental data.

Residue interaction networks of both hArc-NL and hArc-CL were analysed (Fig. 4D) based on the crystal structures, in order to identify residues central to the lobe domain fold. The central residues correspond mainly to buried large hydrophobic residues, which are conserved in the protein family. A Phe residue at the same position of both hArc lobes is has high betweenness in both structures (Fig. 4E).

A structural homology search with hArc-NL and hArc-CL returned, as expected, various capsid proteins from retroviruses, but also other similarly folded helical bundles. The most notable group concerns acyl carrier proteins, which have a similar 4-helix bundle as the Arc NL and CL. For both viral proteins and ACPs, it is noteworthy that the similarly folded proteins often have a sequence identity of less than 5% towards hArc. A selected set of hits is presented in Table 4 and Fig. 5. While it is possible that finding ACPs in the similarity search simply reflects the usefulness of 4-helix bundles for biomolecular interaction modules, to our knowledge, further implications of this structural similarity have not been studied.

**Figure 5.**
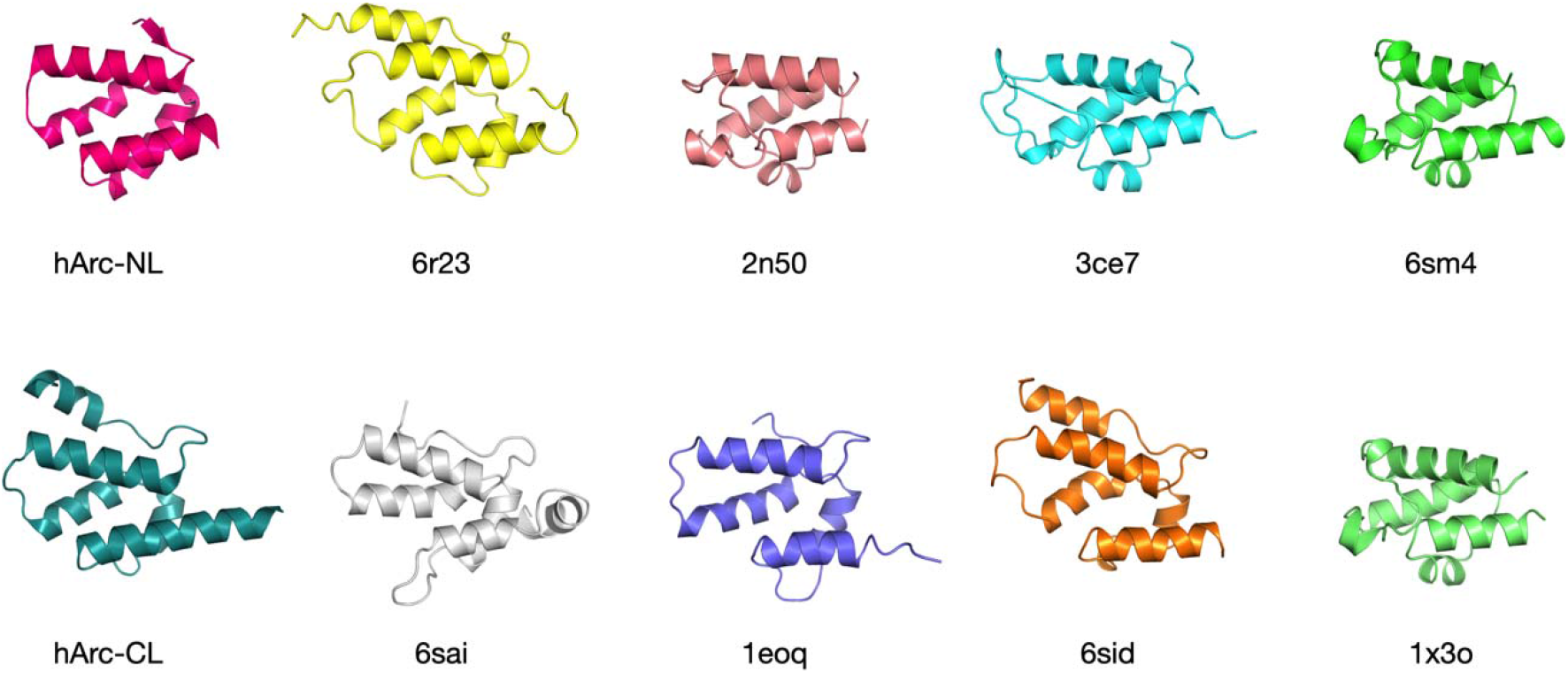
Selected structures with similarity to hArc-NL and hArc-CL. Top row: selected structures similar to hArc-NL. The 3 last ones are different ACPs. Bottom row: selected structures similar to hArc-CL. The last structure is an ACP. See Table 4 for additional details.

**Table 4.** Structural homologues of hArc lobe domains. Shown are selected hits with high similarity from a Salami (40) search with the hArc-NL and hArc-CL structures. Both searches produce a long list of similar structures, with often no sequence identity. Examples of superposed structures are given in Fig. 5.

| Search model | Homologue | PDB entry | Cα RMSD (Å) | Sequence identity (%) | reference |
| --- | --- | --- | --- | --- | --- |
| hArc-NL + pStg(wt) | rat Arc NL + Stg | 4x3h | 0.57 | 100 | (16) |
|  | dArc1 capsid | 6tas | 2.88 | 24 | (44) |
|  | *Toxoplasma* mitochondrial ACP | 3ce7 | 3.44 | 3 | - |
|  | Ty3 capsid | 6r23 | 2.84 | 5 | (65) |
|  | *P. luminescens* ACP | 6sm4 | 3.28 | 2 | (66) |
|  | *E. faecalis* ACP | 2n50 | 3.78 | 3 | (67) |
| hArc-CL | rat Arc CL | 4x3x | 1.56 | 96 | (16) |
|  | dArc1 CL | 6sid | 2.52 | 22 | (45) |
|  | Rous sarcoma virus capsid protein | 1eoq | 2.90 | 18 | (68) |
|  | rat Arc NL | 4x3h | 2.87 | 10 | (16) |
|  | *T. thermophilus* ACP | 1x3o | 4.28 | 5 | - |
|  | Ty3 capsid protein | 6r23 | 2.97 | 12 | (65) |
|  | HML2 C-terminal dimer domain | 6sai | 2.98 | 14 | (69) |

### Arc and the PSD protein network

PSD scaffold proteins belong to several protein families, and they each carry distinct sets of protein interaction domains, which can be used to form multivalent contacts in the tight protein network of the PSD, but also for regulating the PSD molecular assembly in an activity-dependent manner. The latter is important for the participation of the PSD in LTP and LTD. The structure of the PSD varies between excitatory and inhibitory synapses (50).

Different scaffold molecules are concentrated at different depths of the PSD from the postsynaptic membrane (51–53). The distribution of individual proteins can be regulated by synaptic activity (30, 54), which in turn may result in changes in post-translational modifications as well as interaction partners. Some of the PSD proteins, such as the Shanks, have large molecular dimensions and are flexible, possibly enabling them to interact with other proteins at different depths of the PSD assembly (55).

We showed that hArc-NL can bind to multiple sites on GKAP. The function of this interaction remains unknown, but one possible scenario may be that Arc assists in the disassembly/reassembly process of PSD proteins. Large protein complexes in the PSD are formed through a number of relatively weak interactions between multidomain proteins, such as GKAP, PSD95, Shank, and Homer. Could Arc play a role as a rapid regulator and reorganiser of such interactive networks? Putative Arc-binding motifs are present in several components of the PSD assembly (31), but they have not been further characterised.

Phase separation in the PSD is an emerging mechanism of formation of the PSD molecular network (56, 57). The formation of so-called membraneless organelles is central in many biological processes, and phase separation can be induced by properties of individual macromolecules and/or complexes under specific conditions. The structural details of forming a separated phase centered on PSD scaffold proteins, such as GKAP, remain to be elucidated. Recent work using truncated PSD proteins has shed light on protein domains and interactions required for PSD phase separation (58).

GKAP is accumulated at the PSD during synapse inactivity and is removed by synaptic activation due to proteasome-coupled degradation (25). During synapse activation, Arc is rapidly accumulated at the postsynapse, being thereafter degraded by proteasome-coupled degradation (1). An ability of Arc to bind to several proteins in the PSD (31) could allow to disrupt structural interactions between other proteins and therefore “loosen up” the PSD, leading to remodelling of the synapse. Disturbing the interaction between Stg and PSD95 increases AMPAR surface diffusion, preventing AMPAR accumulation at postsynaptic sites (59). AMPAR removal from postsynaptic sites is an effect of Arc expression (13), suggesting that Arc expression and disturbing Stargazin-PSD95 interactions could result in the same outcome. The C terminus of stargazin interacts with the first two PDZ domains of PSD95 (60), while the entire Stg cytoplasmic domain was shown to be involved in the interaction (57).It is the Stg C-terminal tail that interacts with Arc-NL, and it is possible that Arc may disturb the interaction between Stargazin and PSD95.

The PSD95 GK domain interacts with the repeat regions of GKAP (29), and phosphorylation of the GKAP repeats is important for this interaction (61). The repeats of GKAP contain both a putative Arc-binding site and a conserved binding site for PSD95 (Fig. 6A). We have shown that two of these repeats bind to Arc, suggesting that Arc could affect the interaction between PSD95 and GKAP. Structural data at the moment does not necessarily rule out simultaneous binding (Fig. 6B), and Arc oligomerisation could be an additional factor in regulating the interaction. A GKAP mutant incapable of binding PSD95 induced Shank aggregation and degradation in neurons, and Shank and GKAP form aggresomes that are degraded by proteasomes in the absence of PSD95 (62). Due to the multiple possible ligands of Arc, as well as its oligomerisation properties (19, 20), it is possible that a high local concentration of Arc disrupts both the Stargazin-PSD95 as well as the PSD95-GKAP interaction, allowing remodelling of the synapse. Such a function for Arc would explain the burst of Arc protein expression resulting from LTP and rapid degradation, since removal of Arc would be important for the structural stabilisation of the synapse. It would also explain why Arc appears to be involved in both LIP and LTD, as PSD remodelling is required in both situations. Clearly, further experimental studies into the GKAP system, focusing on Arc, are warranted.

**Figure 6.**
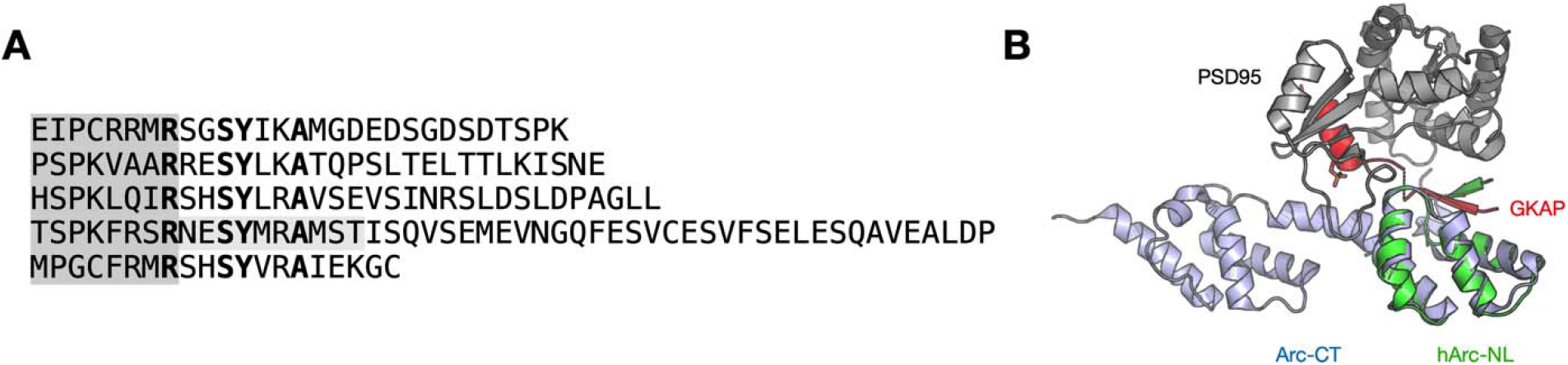
Arc and GKAP. A. Alignment of the five GKAP repeats, including the neighbouring segment that binds to PSD95. Conserved residues in the PSD95-binding site are in bold, and the peptides used in this study are shaded grey. B. 3D model of a ternary complex of Arc, GKAP, and the PSD95 GK domain. The GKAP repeat could sterically bind to both Arc-NL and PSD95 at the same time. Shown are the PSD95-GKAP complex (61), hArc-NL bound to GKAP-R4, and the NMR structure of rat Arc-CT (22).

A further expansion of such a suggested Arc function could include regulation of the actin cytoskeleton, Arc-NL also binds to a peptide from WAVE1 or Wasf1 (16, 23), WAVE1 is an activator of the Arp2/3 complex that is involved in branching of actin filaments in dendritic spines (11). The PSD is connected to actin through cortactin, which binds to Shank *via* its proline-rich domain (63, 64), The interaction of Arc with WAVE1 might be an additional way of affecting the actin cytoskeleton and aiding synaptic remodelling.

### Conclusions

We have shown that hArc-NL binds to specific peptides, with a PxF/Y or P(G)xF/Y motif. Knowing these specific requirements of peptide binding will be helpful when trying to locate Arc binding sites in other proteins and for identifying new Arc binding partners. The peptide binding site of Arc is specific and able to bind to two positions of the GKAP protein, as well as to the phosphorylated and unphosphorylated Stg peptide. For any of the discussed Arc-NL ligands with short linear binding motifs analysed here and in other studies, we currently do not know if additional binding surfaces exist in the corresponding proteins. Such a setting would increase the affinity of the observed interactions, together with possible avidity effects brought about by Arc oligomerisation. The network of interactions between Arc and other PSD proteins is likely to affect the structural arrangement of the postsynaptic density.

## Acknowledgements

We thank the Diamond Light Source (Didcot, UK) for beamtime and support on 103. We also acknowledge EMBL/DESY (Hamburg, Germany), for the provision of experimental facilities at PETRA III and the P13 and P14 beamline staff. Biophysical instrumentation was available through the BiSS core facility at the University of Bergen (Norway). This work was funded through a TOPPFORSK grant from the Research Council of Norway (to CRB)

